# A correlation map of genome-wide DNA methylation patterns between paired human brain and buccal samples

**DOI:** 10.1101/2021.12.09.471962

**Authors:** Yasmine Sommerer, Olena Ohlei, Valerija Dobricic, Derek H. Oakley, Tanja Wesse, Sanaz Sedghpour Sabet, Ilja Demuth, Andre Franke, Bradley T. Hyman, Christina M. Lill, Lars Bertram

## Abstract

Epigenome-wide association studies (EWAS) assessing the link between DNA methylation (DNAm) and phenotypes related to structural brain measures, cognitive function, and neurodegenerative diseases are becoming increasingly more popular. Due to the inaccessibility of brain tissue in humans, several studies use peripheral tissues such as blood, buccal swabs, and saliva as surrogates. To aid the functional interpretation of EWAS findings in such settings, there is a need to assess the correlation of DNAm variability across tissues in the same individuals. In this study, we performed a correlation analysis between DNAm data of a total of n=120 matched *post-mortem* buccal and prefrontal cortex samples. We identified nearly 25,000 (3% of approximately 730,000) cytosine-phosphate-guanine (CpG) sites showing significant (False Discovery Rate *q* < 0.05) correlations between buccal and PFC samples. Correlated CpG sites showed a preponderance to being located in promoter regions and showed a significant enrichment of being determined by genetic factors, i.e. methylation quantitative trait loci (mQTL), based on buccal and dorsolateral prefrontal cortex mQTL databases. Our novel buccal-brain DNAm correlation map will provide a valuable resource for future EWAS using buccal samples for studying DNAm effects on phenotypes relating to the brain. All correlation results are made freely available to the public online.

## Introduction

DNA methylation (DNAm) is an epigenetic mechanism in vertebrate genomes that most often refers to the addition of a methyl-group to cytosine nucleotides within the DNA sequence. Most DNAm in somatic cells occurs in stretches of cytosine-phosphate-guanine (CpG) sites, where they typically, but not always, represent an epigenetic mark of translational repression (1). Owing to the relative technical ease to generate high-resolution DNAm data for thousands of CpG sites simultaneously, it currently represents one of the most frequently studied epigenetic marks. Accordingly, study designs exploiting DNAm profiles on a genome-wide scale – often referred to as epigenome-wide association studies (EWAS) – are becoming increasingly popular. Many EWAS aim to assess the relationship between DNAm patterns and certain brain-related phenotypes (2,3), such as neuropsychiatric traits (4,5), cognitive functions (6,7), and risk for neurodegenerative diseases (8–10), with the goal to better understand the biology and pathophysiology of the traits of interest. However, given that the primary organ of interest, the brain, is typically inaccessible in living individuals, many studies use tissues that are more readily available. While most DNAm studies use blood samples (3,6,7,10), it has been hypothesized that buccal (11) or saliva (12) samples may be more informative for EWAS of psychiatric phenotypes. One key advantage of using peripheral tissues as surrogates is that samples can be obtained from living individuals and do not require *post-mortem* sampling. This also allows for study designs using longitudinal sampling and analysis. However, DNAm patterns are largely cell type- and tissue-dependent (13,14), and therefore it remains unclear how well peripheral DNAm patterns can be used to infer biological processes in the brain.

In the recent past, several attempts have been made to compare DNAm profiles between brain and peripheral tissues within the same individuals. However most studies focused on the comparison of blood and brain tissues (15,16). Only two reports recently compared buccal and brain samples, but sample sizes were very small with 12 and 27 matched sample pairs, respectively (17,18). One additional study used brain, thyroid, and heart tissue samples, each representing one developmental germ layer lineage, from ten individuals to identify so-called correlated regions of systemic interindividual variation (CoRSIV) (19). Interestingly and perhaps not unexpectedly, most of the cited studies report a significant enrichment of methylation quantitative trait loci (mQTL), i.e., DNAm variations that are associated with genetic variants, among CpG sites correlated between tissues or cell-types (13,15–17,19).

Despite this recent progress, there currently remains a significant lack of studies systematically assessing DNAm patterns in paired buccal and brain specimen in sufficiently sized datasets. To close this gap, we here report the results of comprehensive and systematic correlation analyses of genome-wide DNAm patterns in 120 paired prefrontal cortex (PFC) and buccal swab samples. We identified 24,980 significantly correlated CpGs between both tissues and found a significant enrichment of both buccal and dorsolateral prefrontal cortex (DLPFC) mQTLs among the correlated CpG sites. All genome-wide DNAm correlation results are made freely available online (http://www.liga.uniluebeck.de/buccal_brain_correlation_results/), as we anticipate that the buccal-brain DNAm correlation map we generated in this study will provide a valuable resource for the interpretation of EWAS / DNAm studies for brain-related phenotypes.

## Material and Methods

### Human samples

Matched prefrontal cortex (PFC) and buccal samples were obtained in two batches from the neuropathology unit at the Massachusetts Alzheimer’s Disease Research Center (MADRC), Boston, MA, USA. Samples were shipped to our laboratory in two batches: “MADRC-1” and “MADRC-2”, encompassing 48 and 80 matched brain-buccal pairs, respectively. Buccal swabs were obtained from patients with neurodegenerative disease conditions and controls at the time of autopsy following an IRB-approved informed consent with specific inclusion of genetic studies. Consent forms were completed by next-of-kin or other legal representatives as specified by Massachusetts state law. Buccal-Prep Plus DNA Isolation Kit (Isohelix, UK) swabs were utilized to obtain buccal swabs; these were held at -80°C without dehydration until DNA extraction (see below). One hemisphere of each harvested brain was coronally sectioned, flash-frozen on dry ice, cryopreserved at -80°C, and used for subsequent PFC isolation and DNA methylation (DNAm) profiling (see below). The remaining hemisphere was fixed in 10% weight/volume formalin and subjected to detailed neuropathologic evaluation. Detailed descriptions of all MADRC buccal-brain samples used in this study can be found in Supplementary Table 1.

### DNA extraction and processing

DNA was extracted and processed in two laboratory batches according to their shipment charge (i.e. MADRC-1 and MADRC-2; Supplementary Table 1). Importantly, paired brain and buccal samples from the same shipment were processed simultaneously (incl. DNAm profiling, see below). For brain samples, genomic DNA was extracted from approximately 50mg of frozen tissue using the DNeasy Blood & Tissue Kit (Qiagen, Hilden, Germany), while DNA from the buccal swabs was extracted using Buccal-Prep Plus DNA Isolation Kit (Isohelix, UK). All steps in the extraction procedure were conducted according to manufacturer’s instructions. The quantity and the quality of obtained DNA were assessed using a NanoDrop ONE spectrophotometer (Thermo Fisher Scientific, USA).

### EPIC array profiling

DNAm profiling was performed using the “Infinium MethylationEPIC” array (Illumina, Inc., USA), as described previously (8). In brief, experiments were performed on aliquots of DNA extracts diluted to ∼50 ng/µl concentration. Bisulfite conversion of DNA samples was performed using the EZ DNA Methylation kit (Zymo Research, USA), following the alternative incubation conditions for the Illumina Infinium MethylationEPIC Array as recommended by the supplier. After hybridization to the EPIC array, scanning was performed on an iScan instrument (Illumina, Inc.) according to the manufacturer’s instructions (Document#1000000077299v0). DNA samples from both shipment charges (MADRC-1 and MADRC-2) were processed in consecutive laboratory experiments to minimize batch effects. Raw DNAm intensities were determined using the iScan control software (v2.3.0.0; Illumina, Inc.) and exported in .idat format for downstream processing and analysis.

### DNA methylation data processing and quality control

DNAm data from each batch (MADRC-1 and MADRC-2) and tissue (PFC and buccal) were loaded into R and pre-processed separately. DNAm data pre-processing and quality control (QC) was performed using the same procedures as described previously (8) unless noted otherwise. In brief, this entailed using the R (v. 3.6.1) package bigmelon with default settings (20). Samples were excluded when bisulfite conversion efficiency was below 80%. Outliers were removed using the *outlyx* function in bigmelon applying a threshold of 0.15. CpG sites on the X or Y chromosome, or those aligning to SNPs (21) or multiple locations in the genome (22) were removed from the analysis. The final analysis included a total of 120 matched PFC and buccal samples, with 44 samples from MADRC-1, and 76 samples from MADRC-2. Overall, a total of 730,157 QC’ed CpG sites were available in all four datasets and were used for the analyses.

To compare our results with those from the Braun *et al*. study evaluating the correlation of DNAm between buccal and brain samples (17), we downloaded the publicly available .idat files of that study (GEO accession number GSE111165), and loaded and pre-processed them with the R-package ChAMP (23) using default settings unless otherwise noted. Brain and buccal samples were loaded and pre-processed separately. Briefly, DNAm values with a detection *p*-value above 0.01 were set to N/A and CpG sites were completely removed if there were less than 3 beads in more than 5% of the samples, if they were on an X or Y chromosome, or if they aligned to SNPs (21) or multiple locations in the genome (22). Normalization was performed with the BMIQ method. The analysis of this dataset comprised 21 pairs of matched buccal and brain samples, with 740,507 CpG sites. We note that 1,513 (65%) of the 2,367 significantly correlated CpGs according to the Braun *et al*. study (17) were not included in our reanalysis of the dataset as they were removed during QC (Supplementary Table 2). Almost 50% of excluded CpGs (n=931) were removed from our re-analysis of the data due to aligning to or being influenced by SNPs according to Zhou *et al*. (21), while ∼350 CpG sites (23%) were removed due to their location on the X- or Y-chromosome. Despite these differences, we note that the vast majority (713; 83%) of the 854 remaining correlated CpG sites from Braun *et al*. (17) were also significantly correlated in our re-analysis of the Braun *et al*. data after multiple testing adjustment using a False Discovery Rate (FDR) *q*-value threshold of 0.05 (Supplementary Table 2).

### Determination of and correction for DNAm covariates

First, cell-type composition estimates were obtained with the R package EpiDISH (14) for buccal samples and the *estimateCellCounts* function in the R package Minfi (24) for brain samples. Next, to assess the effects of potential confounders on the DNAm data, we used an adaptation of the singular value decomposition (SVD) approach described previously (25). In short, SVD attempts to identify and correct for relevant variables that have a significant impact on genome-wide DNAm patterns and could act as confounders in subsequent analyses. Accordingly, we tested whether variation in cell type composition, bisulfite conversion efficiency, EPIC array ID, diagnosis, extraction date, and position on the EPIC array significantly associated with the variance in DNAm data in our data (as determined by principal component analysis [PCA], see below). These analyses were performed separately for buccal and brain datasets. To this end, we performed a PCA on the DNAm beta values after QC using the R base function *prcomp*. For this PCA, we first generated a subset of uncorrelated CpG sites by dividing the genome into 100kb bins and using one random CpG site from each bin, resulting in 25,746 CpG sites included in each PCA. For the determination of relevant covariates for subsequent analyses, PCs explaining a substantial amount of variance in the DNAm data, as determined by scree plots (MADRC: Supplementary Figure 1; Braun *et al*. data: Supplementary Figure 2) were used. For numerical variables (bisulfite conversion efficiency and cell type composition estimates), a Pearson correlation test between the centred variables and the centred DNAm PCs was calculated with the R base function *cor*.*test*. For categorical variables (extraction date, EPIC array ID, position on the EPIC array, and diagnosis), a one-way ANOVA between the covariates and the DNAm PCs was performed with the R base function *aov*. Effects of variables explaining variance of at least one included DNAm PC with a *p*-value < 0.01 were removed from the DNAm beta values using the *removeBatchEffect* function in the R package limma (26). The results of these analyses, as well as the number of DNAm PC eigenvalues (PCs) included for each dataset, can be found in Supplementary Table 3. The covariate-adjusted DNAm beta values of the two batches of PFC samples and buccal samples were combined in a “brain” and “buccal” data matrix, respectively. Lastly, the batch-defining variable (i.e., indicating either dataset MADRC-1 or MADRC-2) was removed from these combined matrices with the *removeBatchEffect* function in the R package limma (26). All subsequent analyses were performed on these combined DNAm values adjusted for both covariates and batch.

### Identification of CpG sites with correlated DNAm values between paired brain and buccal samples

Spearman rank correlations were calculated for each CpG site in a pair-wise manner across buccal and PFC samples using the R base function *cor*.*test*. The resulting *p*-values were adjusted for multiple testing using the FDR approach with the R base function *p*.*adjust*. FDR *q-*values < 0.05 were considered genome-wide significant in the context of this study. These analyses were performed for the matrices with the two batches (MADRC-1 and MADRC-2) combined, and for each batch separately.

### Identification of mQTLs in buccal samples from an independent dataset

To check for buccal-specific mQTLs, we used an independent in-house dataset with 839 samples ascertained from the Berlin Aging Study II (BASE-II) (27,28). These data are referenced here as “unpublished data”, since a manuscript from our group with more details on this analysis is in preparation. In brief, for this dataset both genome-wide QC’ed DNAm profiles (761,034 CpG-sites) and SNP genotyping data (7,663,257 SNPs) were available for mQTL analysis. DNAm data were derived from buccal-swab samples and generated and processed using the same procedures described above. Genome-wide SNP genotyping data were generated from the same samples using the “Global Screening Array” (GSA) with shared custom content (Illumina, Inc.) using procedures outlined in Hong *et al*. (29). To compute *cis* mQTLs (defined as within ±1 Mb of the CpG site) in this dataset we used the matrix eQTL software (30), which performed an additive linear model with sex, genetic PC 1 to 5, and genotyping batch as covariates. Before association analysis, genome-wide DNAm profiles were adjusted for cell type composition estimates. Only the DNAm and SNP effects that were below a *p*-value of 5.00E-02 were reported. *Cis* mQTLs with FDR *q* < 0.05 were defined as genome-wide significant for this arm of our analyses. Enrichment analyses for mQTLs within CpG sites correlated between PFC and buccal-swab samples were performed with the R base function *chisq*.*test*, using a subset of uncorrelated CpG sites according to 100kb bins, as described above for the PCA.

### Annotation of genomic regions to CpG sites

To assess whether there was a significant enrichment or depletion of CpG sites located in specific genomic regions, we used the annotation from the R package IlluminaHumanMethylationEPICmanifest to assign CpG sites to one of the following genomic regions: 1st exon, 3’ untranslated region (UTR), 5’-UTR, gene body, exon boundary, intergenic region (IGR), the region from transcription start site (TSS) to 200 nucleotides upstream (TSS200), and the region from 200 nucleotides upstream of the TSS to 1,500 nucleotides upstream (TSS1500). Enrichment analyses were performed with the R base function *chisq*.*test*.

### Gene Ontology (GO) analysis

To further characterize the correlated CpG sites, a Gene Ontology (GO) enrichment analysis was performed with the *gometh* function in the R package missMethyl (31) using the significantly correlated (FDR *q* < 0.05) CpGs between PFC and buccal samples. We hypothesized that correlated CpG sites between buccal and brain might show an enrichment for “housekeeping” functions, which would explain the correlated DNAm-values. Nominally significant GO terms were subsequently submitted to the REViGO tool (http://revigo.irb.hr/) (32) to identify and remove redundancy using Resnik’s measure while allowing a terms similarity of 0.7.

## Results

### Spearman rank correlation analysis highlights 24,980 CpG sites showing significant correlation between PFC and buccal samples

Out of the 730,157 CpG sites that were tested, 3% (n=24,980) showed significant Spearman rank correlations of DNAm beta values between paired PFC and buccal samples after adjustment for multiple testing (FDR *q* < 0.05). Most of the significantly correlated CpG sites (n=24,636; 99%) had a positive Spearman rank correlation coefficient (Figure 1), which means that DNAm patterns were consistent in both tissues. The remainder (n=344; 1%) showed negative correlations, meaning that the effect direction was opposite in buccal and brain samples. Furthermore, correlated CpG sites were evenly distributed across the genome and showed no obvious preponderance for any significant genomic location (Supplementary Figure 3). Lastly, the majority of significantly correlated CpG sites was also correlated between PFC and buccal samples at *p* < 0.05 in analyses that were performed for each batch (“MADRC-1” and “MADRC-2”) separately (Supplementary Figure 4). These latter computations were performed to check for spurious correlations that may have resulted from undetected confounding after merging both laboratory batches.

**Figure 1:**
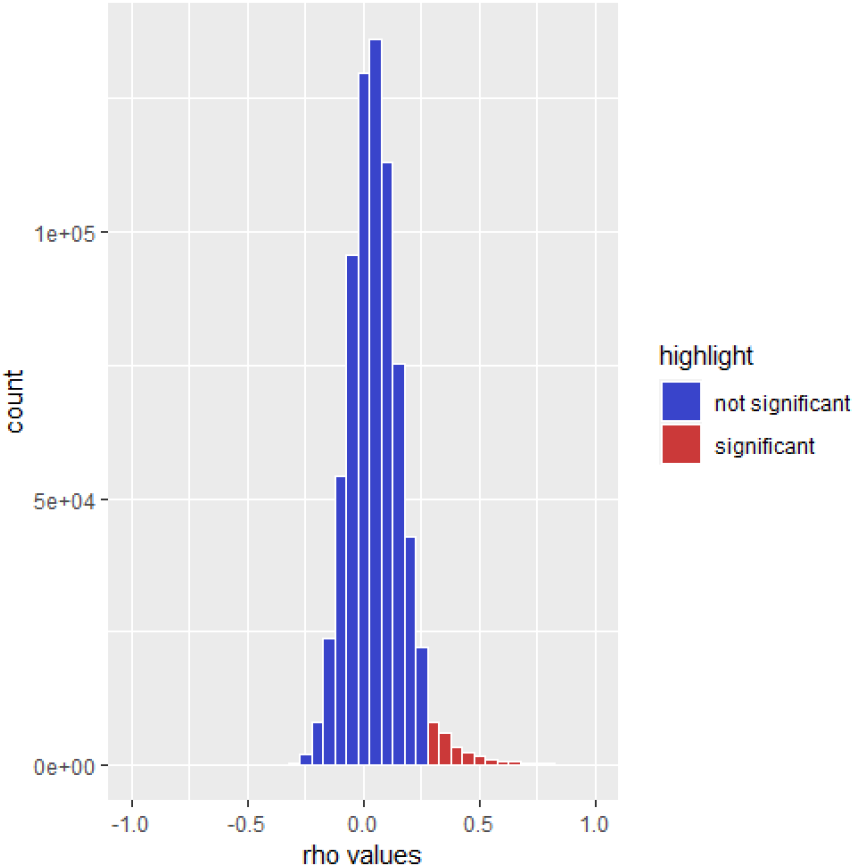
Histogram of Spearman rank correlation coefficients. Significantly correlated CpG sites between PFC and buccal samples after multiple testing adjustment (FDR q < 0.05) are highlighted in red.

### Correlated CpG sites are enriched for both buccal and brain mQTLs

We next looked up our ∼25K correlated brain-buccal CpGs in two mQTL databases: Firstly, in an mQTL database (https://eqtl.brainseq.org/WGBS_meQTL/) for dorsolateral prefrontal cortex (DLPFC) (33) mQTLs based on 165 samples, and secondly, in an in-house mQTL database for buccal samples that we generated in 839 individuals analysed as part of the Berlin Ageing Study II (BASE-II; unpublished data). The look-up resulted in an enrichment of *cis* mQTLs in the fraction of CpG sites that were correlated between PFC and buccal samples when compared to all analysed CpGs (*p* = 2.45E-09 for DLPFC mQTLs and *p* = 1.13E-81 for buccal mQTLs using a Chi-squared test, Figure 2). Overall, 22,055 CpG sites (88%) that were significantly correlated between PFC and buccal samples in our dataset were identified in mQTL analyses in either buccals (n=20,940) or DLPFC (n=12,888). Of these, 11,773 CpG sites were identified in mQTL analyses in both tissues.

**Figure 2:**
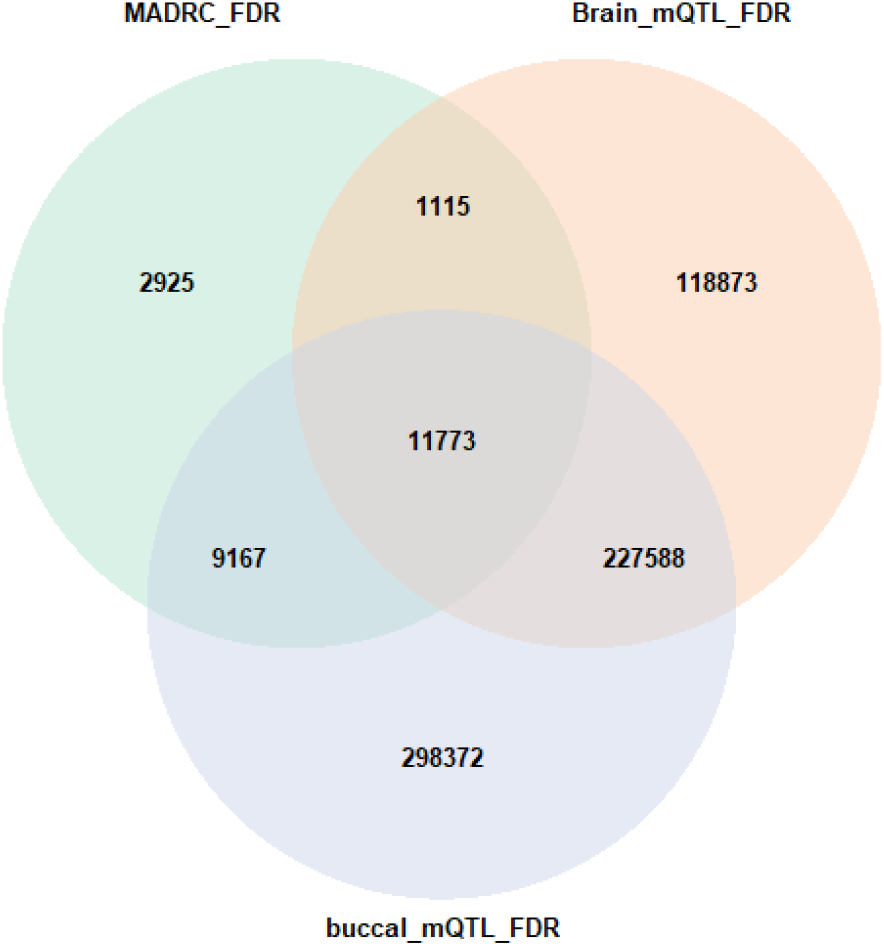
Venn diagram showing significantly correlated CpG sites between PFC and buccal samples after multiple testing correction in the MADRC dataset (n=24,980); Overlap of significantly correlated CpG sites between PFC and buccal samples (FDR q < 0.05; green) with mQTLs according to the DLPFC mQTL database (FDR q < 0.05; orange) (33) and the buccal mQTL database generated from an independent dataset ([unpublished data], FDR q < 0.05; violet).

### Correlated CpG sites show good concordance with results from published datasets using matched brain and peripheral tissues

We compared our results to those of two previous publications evaluating the correspondence of DNAm between tissues (17,19). The first study by Braun *et al*. (17) correlated DNAm-values between brain tissues obtained during a surgery of epilepsy patients and buccal samples from the same individuals. Overall, they identified 2,367 CpGs (i.e. 0.29% of all tested) as significantly correlated (17). To increase comparability between our and the Braun *et al*. dataset (17), we reprocessed the DNAm raw data from that study including covariate adjustment and Spearman rank correlations as applied to the MADRC datasets. Our re-analysis of the Braun *et al*. data (for details of the analysis see methods section) resulted in 27,796 CpGs showing significant (FDR *q* < 0.05) correlation between buccal and brain tissue. While this number is comparable to the 24,980 correlated probes identified in the analyses of the MADRC data, it still represents a nearly 10-fold difference as compared to the numbers originally published by Braun *et al*. (17). This (stark) difference can likely be attributed to a more stringent multiple testing correction procedure (i.e. Bonferroni [Braun *et al*.] vs. FDR [here]) and differences in data processing and analysis strategies (Methods). Overall, a total of 5,918 (24%) of the 24,980 correlated CpGs in our data also represented correlated CpGs in the Braun *et al*. data.

The second dataset used for comparison was recently published by Gunasekara *et al*. (19) who identified 9,926 significant CoRSIVs across three different tissues (brain, thyroid, and heart) from 10 matched individuals. A total of 1,311 (13%) of all CoRSIVs also had at least one CpG probe on the EPIC array, with some regions being represented by more than one CpG site. This resulted in a total of 1,684 individual CpG sites in the MADRC analysis that were located in a CoRSIV and could be used for comparison. A total of 897 CpG sites (53%) showed a significant correlation at FDR *q* < 0.05 in our data, too. For a depiction of the three-way comparisons of the Braun *et al*., Gunasekara *et al*., and our MADRC data, see the Venn diagram in Figure 3.

**Figure 3:**
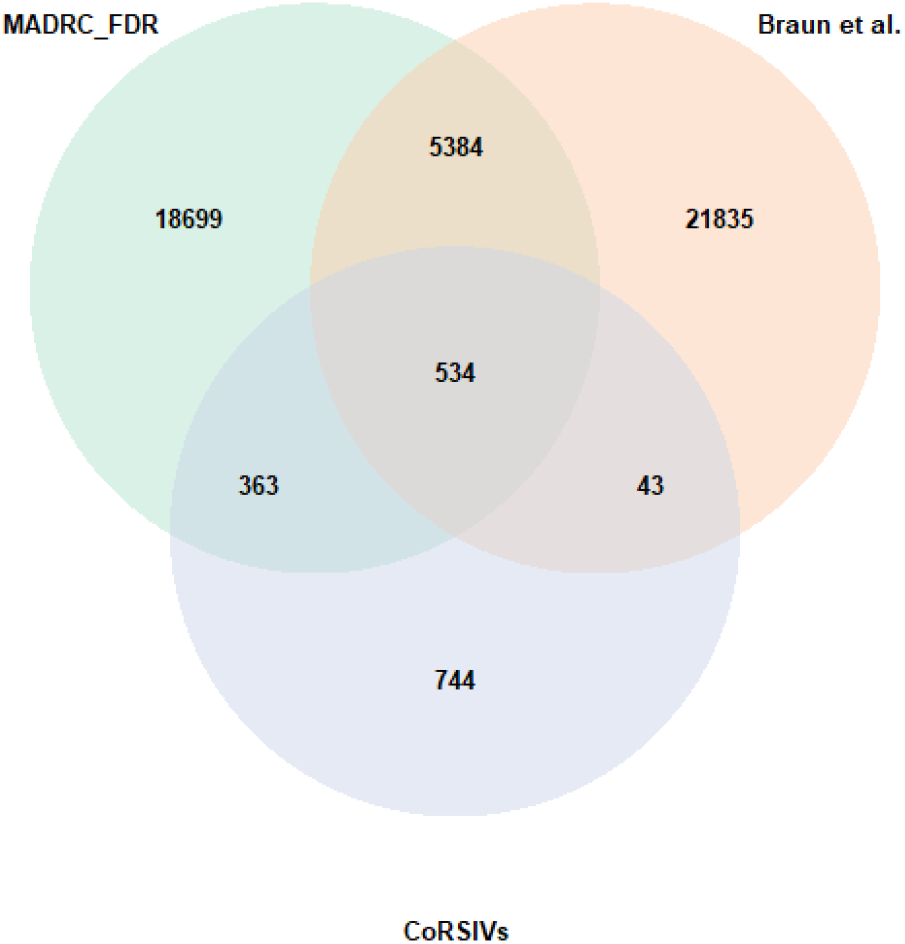
Venn diagram comparing significantly correlated CpG sites between buccal and brain samples in our analysis (MADRC; green), the analysis using Braun et al. (17) data (orange), and CoRSIVs (violet) (19).

### Genomic location of correlated CpG sites shows an enrichment in gene promoters and depletion in gene bodies

Next, we assessed whether there was an enrichment or depletion of correlated CpG sites in specific genomic regions in comparison to all CpGs. We noted a statistically significant change of the genomic region distribution within the CpG sites that were correlated between PFC and buccal sample DNAm in the MADRC dataset compared to the expected distribution in our data (Chi-squared test with *p* < 2.20E-16, Figure 4). Specifically, we observed an enrichment of CpG sites located in the 1st exon, intergenic regions, regions 1,500 nucleotides upstream of transcription start sites, and regions 200 nucleotides upstream of transcription start sites, and a depletion in the 3’-UTR, 5’-UTR, gene bodies, and exon boundaries within the correlated CpG sites (Figure 4). The enrichment in IGRs and depletion in gene bodies is in line with previous observations made for CoRSIVs (19).

**Figure 4:**
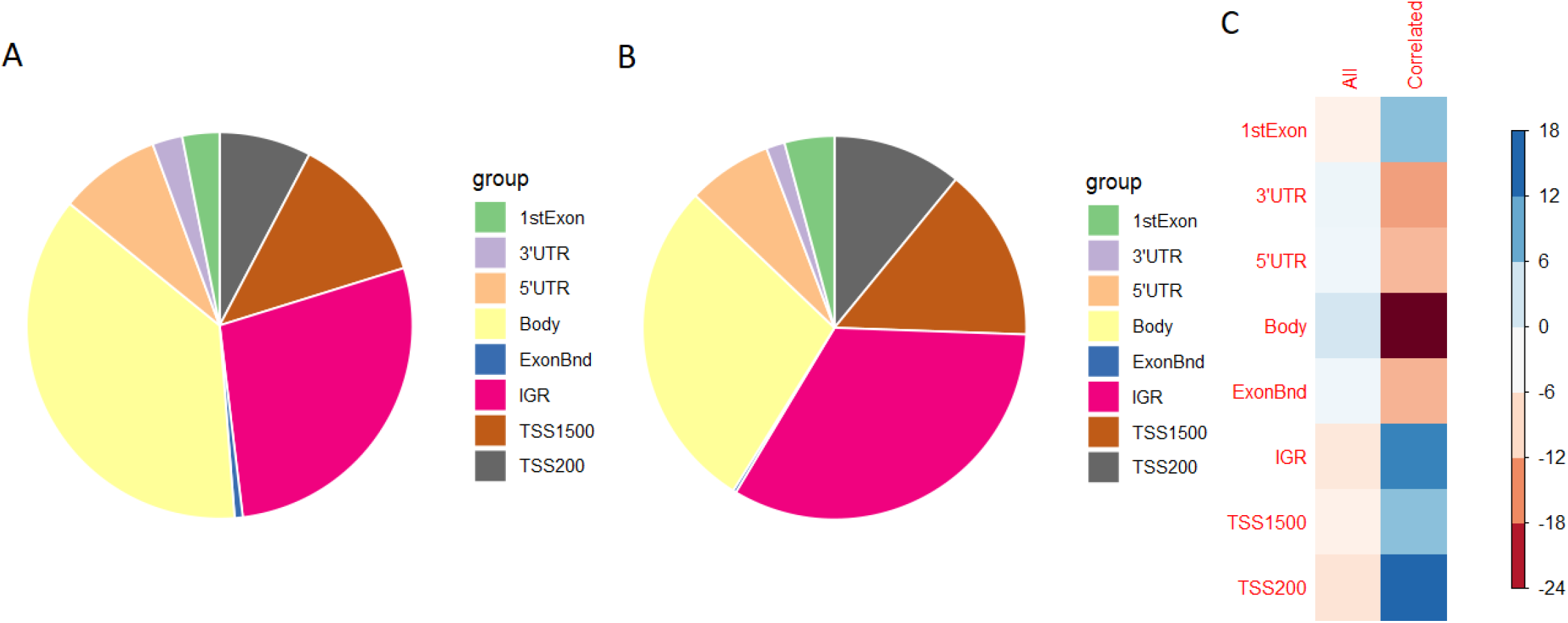
Enrichment of genomic regions within the significantly correlated CpG between buccal and brain samples in the MADRC dataset; Panel A: Distribution of genomic regions in all tested CpG sites on the EPIC array; B: Distribution of genomic regions in significantly correlated (FDR q < 0.05) CpG sites; Panel C: Pearson’s residuals of the Chi-squared test, blue indicates an enrichment and red a depletion of the respective genomic region

### GO analysis of genes annotated to correlated CpG sites highlights cellular functions relating to major histocompatibility complex (MHC)

Next, we aimed to assess whether the correlated buccal-brain CpG sites fell into specific functional pathways and tested for an enrichment of specific gene ontology (GO) terms. Upon including all 24,980 significantly correlated (FDR *q* < 0.05) CpG sites, this analysis revealed only one statistically significant GO term (GO:0007156, “homophilic cell adhesion via plasma membrane adhesion molecules”, FDR *q*=0.04). In order to remove redundant terms, all nominally significant GO terms (*p* < 0.05) were submitted to REViGO, which resulted in 246 nominally significant GO terms (Supplementary Table 4). Among the top GO terms showing at least a nominally significant enrichments were many terms related to “housekeeping” functions, such as peptide antigen binding, MHC protein complex, ion binding, cell-cell adhesion via plasma-membrane adhesion molecules, transferase activity, manganese ion binding, and nucleoside-triphosphatase regulator activity (Supplementary Table 4). In general, this is in line with the GO results presented in the Gunasekara *et al*. (19) publication (using GO terms associated with the CoRSIVs). One noteworthy overlapping annotation was observed with the “MHC protein complex” likely highlighting the central role of MHC-mediated immune response in both tissues.

### Look-up of Alzheimer’s disease EWAS results as an example of application for the buccal-brain correlation map

As a first “practical” application of our buccal-brain correlation results, we looked up top-hits from the hitherto largest EWAS across several brain regions (9) and for the entorhinal cortex (EC) for Alzheimer’s disease (AD) (8). This look-up revealed that five of the CpGs that were previously highlighted in the context of AD also showed significant correlations at FDR *q* < 0.05 and all correlations were positive (cg05030077 [MLST8/chr16:2255199], cg05923197 [BMP4/chr14:54418804], cg23950714 [DOK3/chr5:176935364], cg04252044 [chr3:188664747], cg24569831 [RGMA/chr15:93617168], Supplementary Table 5). Probe cg05923197 is also an mQTL according to the DLPFC mQTL database (33) and all five of them were identified as mQTLs in our buccal mQTL database (unpublished results). Furthermore, lowering the significance threshold to correlations significant at a nominal *p*-value of 0.05 increased this number to 29 CpGs showing association in one of the AD brain EWAS. Among these 29, 8 CpG sites were reported to be mQTLs in the DLPFC and 28 in the buccal databases. Interestingly, all but two showed positive correlations in DNAm beta values between brain and buccal tissues (Supplementary Table 5). Based on these data, we predict that all 29 overlapping CpGs should show at least some degree of association evidence in AD EWAS performed in buccal samples. Their potential for representing early biomarkers of AD should be the focus of future work.

## Discussion

In this study, we generated high-resolution genome-wide DNAm profiles from brain and buccal samples collected *post mortem* from the same individuals at the same timepoint. Comparing CpG sites across both tissues revealed ∼25K sites showing significant correlations in DNAm levels. This is in line with a previous study assessing the correlation of DNAm between brain and blood samples, which reported moderate to strong correlations for 1% to 6% of CpG sites (16). Correlated CpGs showed an enrichment for being regulated by mQTLs using both buccal and brain databases, which is in line with results from previous publications (13,15–17,19). In terms of physical location, correlated CpGs showed a significant enrichment in promoter regions and a significant depletion in gene bodies. A GO enrichment analysis highlighted terms related to molecular “house-keeping” functions, including several significant GO terms linked to the MHC protein complex, confirming previous findings (19). To our knowledge, our study has generated the largest buccal-brain DNAm correlation map available to date and will hopefully prove to be a valuable resource for the interpretation of EWAS for brain-related phenotypes. To this end, we made all of our genome-wide DNAm correlation results freely available online (http://www.liga.uniluebeck.de/buccal_brain_correlation_results/).

Comparing our correlation results with those from a previous publication from Braun *et al*. (17) using matched brain and buccal samples resulted in an overall good correspondence in results (Supplementary Table 2). Differences in findings across both studies can likely be attributed to differences in clinical diagnoses of included individuals, as well as differences in technical aspects such as study design and QC procedures. For instance, the brain samples from Braun *et al*. (17) were obtained from living individuals during epilepsy surgery, whereas the MADRC samples were collected *post mortem* and included individuals with different neurodegenerative diseases (Supplementary Table 1; see below). As a result, the brain samples from the Braun *et al*. study (17) were from many different brain regions, whereas the MADRC brain samples were all obtained from the PFC. In addition, in the MADRC dataset the extraction of brain and buccal samples was performed at the same time, i.e. at the time of autopsy, whereas brain and buccal samples in the Braun *et al*. study (17) were not always collected at the same time point (time range 0-638 days). The second comparison with published data was performed with the CoRSIVS from Gunasekara *et al*. (19). Although the CoRSIV assessments from that study and buccal-brain correlation analyses performed here differed in many important aspects, the overlap between both analyses was more than 50% of all CpGs on the EPIC array that are located in CoRSIVs. We suggest that this number, rather than the 21% overlap with Braun *et al*., be considered as the lower bound of “true” correlations in DNAm patterns in human buccal vs. brain samples.

The strengths of our study are its pair-wise design (i.e. simultaneous collection and analysis of all paired buccal and brain samples), the use of the most current DNAm microarray (i.e. the EPIC array with ∼730K CpG probes available for analyses as opposed to the predecessor array with 450K CpGs), our stringent QC and data processing procedures (e.g. to eliminate bias due to undetected confounding by certain biological or technical variables), and the comparatively large size of our sample (i.e. n=120 here versus n=21 in Braun *et al*. (17) and n=10 in Gunasekara *et al*. (19)). Furthermore, we make use of a novel and hitherto unpublished buccal tissue mQTL database from our group, allowing to determine the impact of genetics in the greatest detail possible. Despite these strengths, our study is also subject to several limitations. First and probably most importantly, all DNAm profiles obtained from this study are from bulk tissue samples. While all DNAm data were corrected for cell type composition estimated from current reference panels, it cannot be excluded that undetected difference in cell type composition across samples has created a bias in results. Conceptionally, however, this bias (if it existed) should have increased the number of false-negative findings but would not invalidate our findings, i.e. it would result in a bias towards the null. Only studies applying single-cell sequencing could shed more light on the impact of cell-type specific differences in DNAm profiles. Second, we note that the majority of individuals used in this study were not “healthy controls” but had received some type of neuropathological diagnosis, mostly due to the presence of some neurodegenerative disorder (Supplementary Table 1). Since these underlying disease conditions likely had in impact on DNAm patterns in the brain and treatment regimens may have affected methylation in the brain and elsewhere, it cannot be excluded that the buccal-brain correlation results reported here were at least partially influenced by diagnosis status. In a similar vein, all subjects included here were relatively old, with a mean age of ∼72 years. It is well known that DNAm patterns change over time and are different in aged vs. non-aged individuals (34–36), so use of our buccal-brain correlation map may be less informative for EWAS of younger individuals. Third, as described above, both types of sampled biospecimen (i.e. brain and buccal swabs) were collected *post mortem*, i.e. with a specific and varying time interval (*post-mortem* interval; PMI) between death and sampling (Supplementary Table 1). It is difficult to predict whether and how DNAm patterns were affected by differing PMIs across individuals and tissues. However, evidence from previous work suggests that DNAm appears to be a rather stable epigenetic marker under varying conditions in brain samples collected *post-mortem* (37). Lastly, despite stringent QC of the DNAm, including adjustment for covariates that may have had a substantial influence on DNAm (Methods) we cannot exclude that some correlations reported in this study are the result of some unknown and undetected confounding. However, given the pair-wise design of our study where we used matched buccal and brain samples from the same individuals, it appears unlikely that undetected confounding has led to a substantial and systematic inflation of our results. Notwithstanding, future studies are needed to verify and replicate the findings we present here, ideally using single-cell DNAm assessments in sufficiently sized samples.

In summary, our study on genome-wide DNAm patterns in paired buccal and brain samples highlighted ∼25K sites showing significant correlations in DNAm levels across both tissues. To our knowledge, our study is the largest buccal-brain DNAm correlation map available to date and will hopefully prove to be a valuable resource for the interpretation of EWAS/DNAm studies for brain-related phenotypes. To this end, we made all of our genome-wide DNAm correlation results freely available online (http://www.liga.uniluebeck.de/buccal_brain_correlation_results/).

## Declarations

### Ethics approval and consent to participate

#### MADRC

Sample extraction at the time of autopsy was done according to an IRB-approved informed consent with specific inclusion of genetic studies. Consent forms were completed by next-of-kin or other legal representatives as specified by Massachusetts state law.

#### BASE-II

All participants gave written informed consent. The medical assessments at baseline and follow-up were conducted in accordance with the Declaration of Helsinki and approved by the Ethics Committee of the Charité – Universitätsmedizin Berlin (approval numbers EA2/029/09 and EA2/144/16).

### Consent for publication

Not applicable.

### Availability of data and materials

All results pertaining to the buccal-brain DNAm correlation analyses are freely available via http://www.liga.uni-luebeck.de/buccal_brain_correlation_results/. The DNAm raw data generated and used for the analyses described in this manuscript are not publicly available due to data sharing restrictions.

### Competing Interest Statement

BTH has a family member who works at Novartis, and owns stock in Novartis; he serves on the SAB of Dewpoint and owns stock. He serves on a scientific advisory board or is a consultant for AbbVie, Avrobio, Axon, Biogen, BMS Cell Signaling, Genentech, Ionis, PPF, Novartis, Seer, Takeda, the US Dept of Justice, Vigil, Voyager. His laboratory is supported by Sponsored research agreements with Abbvie, F Prime, and research grants from the National Institutes of Health, Cure Alzheimer’s Fund, Tau Consortium, and the JPB Foundation. The other authors declare no conflict of interest.

### Funding

This work was supported by the Cure Alzheimer’s Fund (as part of the “CIRCUITS” consortium) to LB, Deutsche Forschungsgemeinschaft (DFG) and the National Science Foundation China (NSFC) as part of the Joint Sino-German research project (“MiRNet-AD”, #391523883) to LB, and by the EU Horizon 2020 Fund (as part of the “Lifebrain” consortium, #732592) to LB. The BASE-II research project (Co-PIs: Lars Bertram, Ilja Demuth, Denis Gerstorf, Ulman Lindenberger, Graham Pawelec, Elisabeth Steinhagen-Thiessen, and Gert G. Wagner) is supported by the German Federal Ministry of Education and Research (Bundesministerium für Bildung und Forschung, BMBF) under grant numbers #16SV5536K, #16SV5537, #16SV5538, #16SV5837, #01UW0808, 01GL1716A, and 01GL1716B. The Massachusetts Alzheimer’s Disease Research Center is supported by the National Institute on Aging NIA (Grant P30AG062421).

### Author contributions

Design of the study, supervision, and acquisition of funding: LB. Ascertainment of buccal tissue: DHO, BTH, ID. Ascertainment of brain tissue and neuropathological examinations: DHO, BTH. Handling of tissue samples and generation of molecular data: VD, TW, SSS, AF. Data processing and statistical analyses: YS, OO, CML. First draft of the manuscript: YS, LB. Critical revision and final version of manuscript: all authors.

## Acknowledgements

We acknowledge the high-performance compute environment (“OmicsCluster”) at University of Lübeck where most data processing and analysis steps of this study were run.

## Supplementary Figures

**S1:**
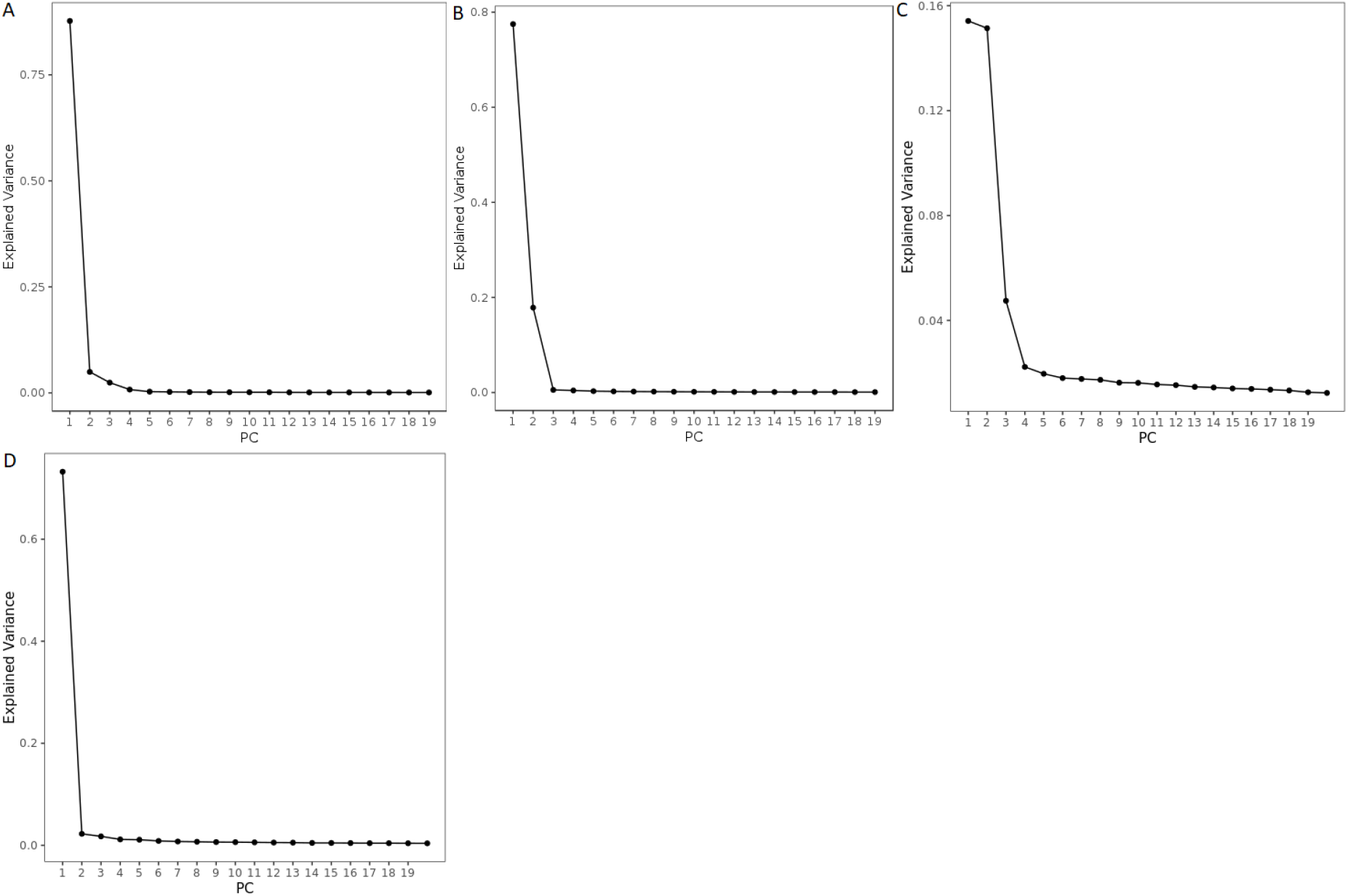
Scree plots of PCA of normalized DNAm beta values in MADRC-1 brain (A, DNAm PC1 was used for analyses), buccal (B, DNAm PC 1 and 2 were used for analyses), MADRC-2 brain (C, DNAm PC1 to 3 were used for analyses), buccal (D, DNAm PC 1 was used for analyses)

**S2:**
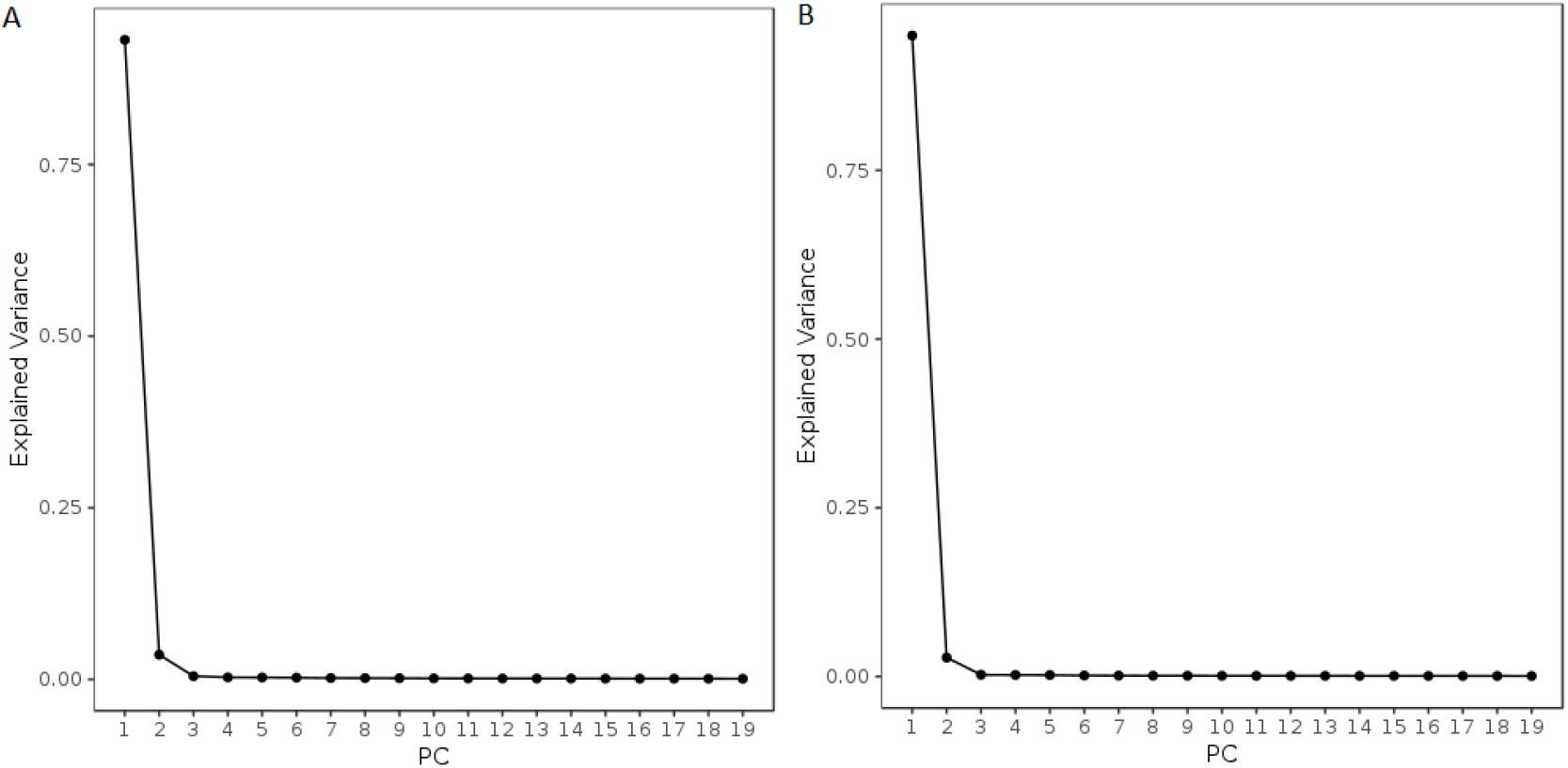
Scree plots of PCA of normalized DNAm beta values in the Braun et al. data for brain (A, DNAm PC 1 was used for analyses) and buccal (B, DNAm PC 1 was used for analyses) samples

**S3:**
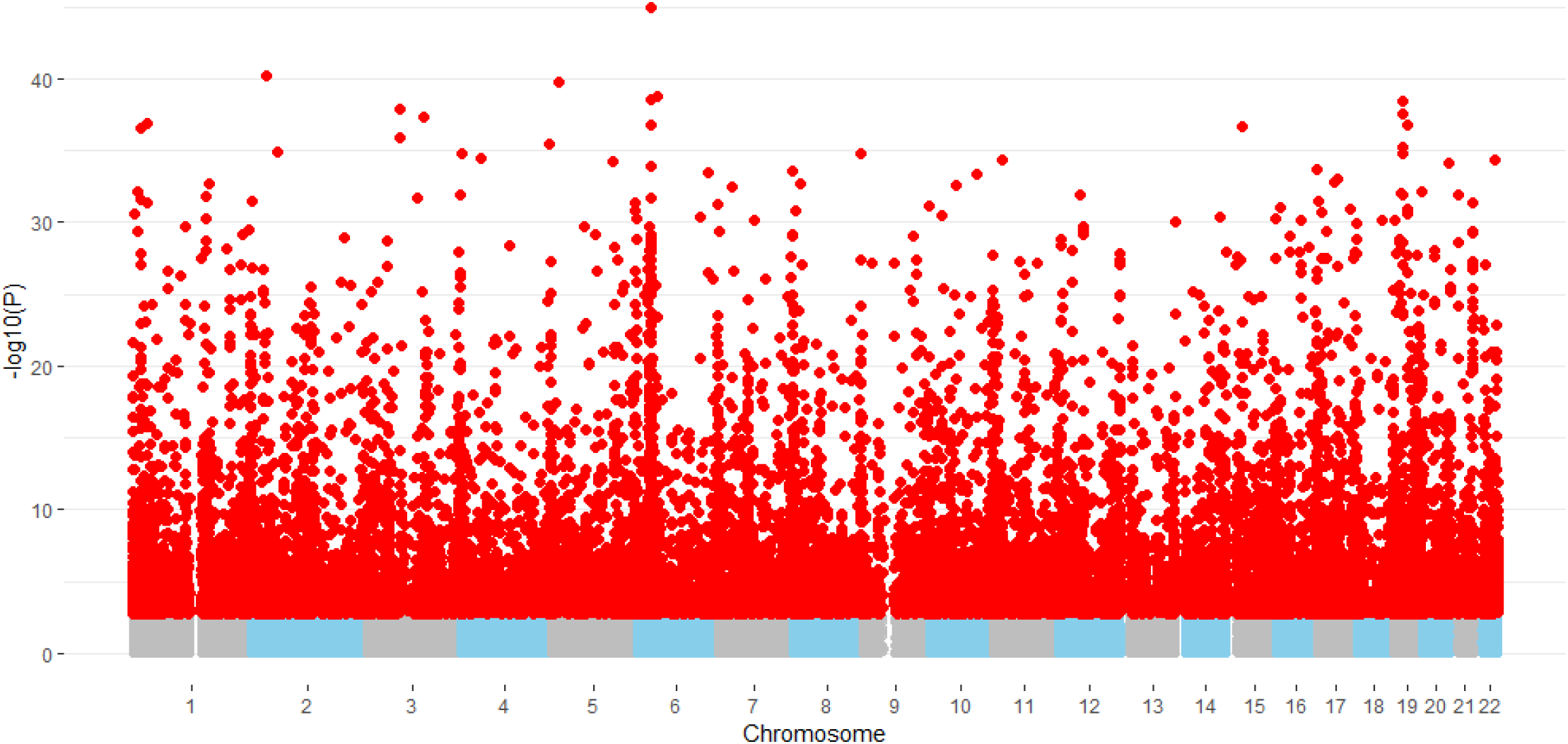
Manhattan plot of the Spearman rank correlation test results. P-values are plotted and significantly correlated CpG sites after FDR multiple testing correction (q < 0.05) are highlighted in red.

**S4:**
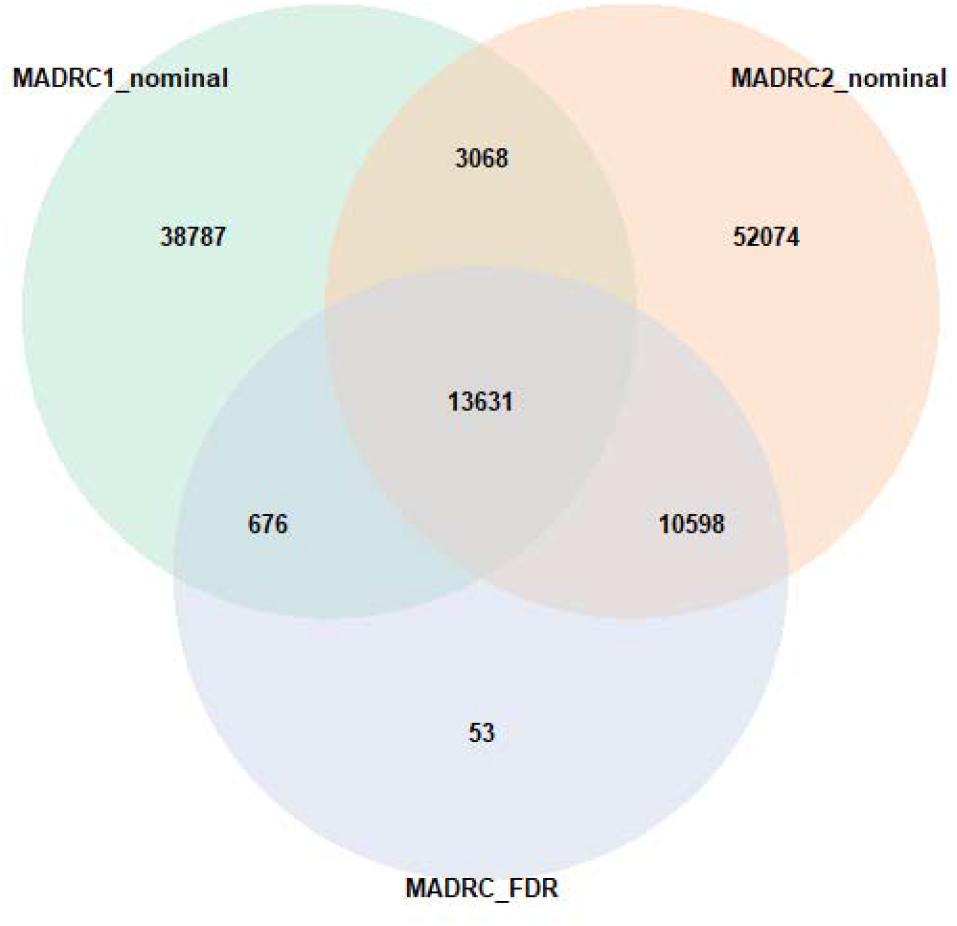
Venn diagram displaying the overlap between CpG probes that are correlated between PFC and buccal samples with a nominal p-value below 0.05 in the analyses in each individual batch (green and red), and the significantly correlated CpG sites in the combined analysis (purple).

## Notes

http://www.liga.uni-luebeck.de/buccal_brain_correlation_results/

